# eIF2α phosphorylation evokes dystonia-like movements with D2-receptor and cholinergic origin and abnormal neuronal connectivity

**DOI:** 10.1101/2024.05.14.594240

**Authors:** Sara A. Lewis, Jacob Forstrom, Jennifer Tavani, Robert Schafer, Zach Tiede, Sergio R. Padilla-Lopez, Michael C. Kruer

## Abstract

Dystonia is the 3^rd^ most common movement disorder. Dystonia is acquired through either injury or genetic mutations, with poorly understood molecular and cellular mechanisms. Eukaryotic initiation factor alpha (eIF2α) controls cell state including neuronal plasticity via protein translation control and expression of ATF4. Dysregulated eIF2α phosphorylation (eIF2α-P) occurs in dystonia patients and models including DYT1, but the consequences are unknown. We increased/decreased eIF2α-P and tested motor control and neuronal properties in a Drosophila model. Bidirectionally altering eIF2α-P produced dystonia-like abnormal posturing and dyskinetic movements in flies. These movements were also observed with expression of the *DYT1* risk allele. We identified cholinergic and D2-receptor neuroanatomical origins of these dyskinetic movements caused by genetic manipulations to dystonia molecular candidates eIF2α-P, ATF4, or DYT1, with evidence for decreased cholinergic release. *In vivo*, increased and decreased eIF2α-P increase synaptic connectivity at the NMJ with increased terminal size and bouton synaptic release sites. Long-term treatment of elevated eIF2α-P with ISRIB restored adult longevity, but not performance in a motor assay. Disrupted eIF2α-P signaling may alter neuronal connectivity, change synaptic release, and drive motor circuit changes in dystonia.

## Introduction

Eukaryotic initiation factor (eIF2) is the protein translation complex that brings initiator tRNA, mRNA, and the ribosome together for protein translation under basal conditions ^1^. Phosphorylation of the eIF2 alpha subunit (eIF2α-P) regulates protein translation as a finely controlled mechanism for responding to diverse cellular states and is part of the integrated stress response (ISR) ^2^. eIF2α can be phosphorylated by the kinase GCN2 (which senses essential amino acid deprivation) ^3^, PKR (which is activated by virus-associated dsRNA) ^4^, and PEK (which senses ER stress from unfolded protein accumulation) ^5^. In response to these stressors, phosphorylation selectively regulates mRNA translation by decreasing 5’UTR uORF sequence affinity for the ribosome ^6^ resulting in a global decrease in protein synthesis. Transcripts that are inhibited by uORF are preferentially upregulated, such as the transcription factor ATF4; ATF4 regulates gene expression of factors for apoptosis, autophagy, ^7^ as well as neuron plasticity ^8^. High levels of ATF4 initiate negative feedback to reset the pathway by increasing expression of the growth arrest and DNA damage-inducible protein (GADD34/PPP1R15), the regulatory subunit 15A for the phosphatase removes the phosphate group from eIF2α ^9^. Sustained phosphorylation of eIF2α can result in significant downregulation of the rate of protein synthesis ^10^ and disproportionate ATF4-mediated responses ^7^.

Separate from stress, eIF2α-P and ATF4 are increasingly recognized for neuron-specific states including process outgrowth, plasticity, neuron excitability, and synaptic release. ATF4 regulates cellular responses to growth factors such as BDNF ^11^ and guidance cues such as Sema3a during axon pathfinding ^12^ without initiating other aspects of the ISR. ATF4 is also a known inhibitor of CREB-mediated neuronal plasticity and is required to be degraded for induction of LTP ^13^. ATF4 positively regulates trafficking of metabotropic GABA_B_ receptors via localization to postsynaptic membrane with ATF4 knockdown leading to neuronal hyperexcitability ^14^. Further, pharmaceutical induction of eIF2α-P decreases expression of exocytosis components and subsequent release of synaptic-like vesicles, suggesting this pathway could also be crucial for regulating synaptic release ^15^.

Dystonia is the third most common movement disorder in humans, causing abnormal, unwanted muscle activation that results in twisting and/or posturing movements that can be focal or generalized. Dystonia may respond to medications, injection therapies (botulinum toxins or phenol), or deep brain stimulation but it is often painful for affected individuals and refractory to medical treatment. Although advances have been made in understanding dystonia at the genetic and circuit levels utilizing models of *DYT1*, progress in the field has been hampered by limited knowledge of the molecular and cellular substrate of dystonia that connects genetic risk factors and altered brain circuit connectivity ^16^.

Altered functional connectivity between brain regions including sensory-motor cortex, thalamus, cerebellum, and basal ganglia has been observed in dystonia ^17,18^. The striatum of the basal ganglia is a potential anatomical location of these alterations due to its role regulating motor control via the direct vs indirect motor pathways ^19^. Candidate underlying neuronal mechanisms of this altered connectivity include changed wiring between neurons, neuronal hyperexcitability, or changes in synaptic strength. Synaptic connectivity changes have been observed as increased dendrite arbor size of the medium spiny neurons, the GABAergic interneurons of the striatum, in *DYT1* knock-in mouse model ^20^. Alterations to neuron excitability has been observed as decreased firing of striatal neurons of the indirect pathway in a Paroxysmal Dyskinesia (PNKD) mouse model during dyskinesia bouts ^21^. Increased synaptic connectivity from excessive long-term potentiation (LTP) and impaired long-term depression (LTD) have been observed in several dystonia models ^22,23^. Therefore, multiple types of neuronal properties could be altered in dystonia, although how they arise and whether they co-occur needs further investigation.

Striatal cholinergic neurons and neurons expressing D2 receptors in the indirect motor pathway are potential cellular origins of dystonia, although the exact neurobiology of dystonia is unknown. Anticholinergic medications can treat dyskinetic symptoms and both levodopa and dopamine agonists can both treat dyskinetic movements and induce dyskinetic movements with long-term use ^24,25^. In a rat model of term-born hypoxic-ischemic encephalopathy with dystonic movements, the number of striatal cholinergic interneuron increased following neonatal brain injury while the number of parvalbumin-positive GABAergic striatal interneurons are unchanged ^26^. Chronic stimulation of striatal interneuron cholinergic neurons leads to the development of dystonia-like movements in a rodent model (bioRxiv doi: 10.1101/2023.07.19.549778). Striatal cholinergic neurons are both increased in size ^20^ and more excitable ^27^ in rodent models of *DYT1* dystonia. Choline can both increase and decrease dopamine neuron activity and subsequent dopamine release in the striatum through nicotinic ^28^ and M4 metabotropic ^29^ receptors, respectively. There is reduced D2 receptor binding in the striatum detected during radioimaging in isolated idiopathic dystonias ^30^. Together, this suggests a model of dystonia where choline positively regulates dopamine levels, inhibiting D2-receptor expressing neurons of the indirect motor pathway, and thereby inhibiting unwanted movements.

The genetic landscape of dystonia is diverse, but eIF2α is emerging as a potential point of convergence. The best studied genetic form of dystonia, *DYT1*, has altered eIF2α-P in patient tissues, cellular models, and animal models ^31^. ATF4 expression was poorly upregulated in response to stress in *DYT1* patient cells compared to controls ^32^ and was reduced in postmortem brain tissue despite increased eIF2α-P ^33^. Phosphorylation of eIF2α was upregulated despite decreases activity of the activating kinases PKR and PERK in a *DYT1^ΔE^* risk allele expressing rat model ^33^. Decreased expression of eIF2α-P regulators, including ATF4, was also observed in a *DYT6* rat model ^23^. The fly ortholog of the dystonia-associated gene *AGAP1* has chronically elevated eIF2α-P with the inability to increase phosphorylation in response to misfolded proteins and starvation stress ^34^. Finally, genes regulating eIF2α-P and translation initiation/elongation factor genes represent monogenetic causes of dystonia including *PKR/EIF2AK2* ^35^ and *EIF2AK1* ^36^, *PRKRA/DYT16* ^37^, *EIF4A2* ^38^, and EEF1A2 ^39^.

Attempts to correct eIF2α-P defects in these models suggests that this is a candidate molecular mechanism for dystonia pathophysiology. Increasing eIF2α-P improved mislocalization phenotypes in *DYT1* patient cells and defective LTD in *DYT1* mice ^32^. Suppressing physiologic eIF2α-P activation during the initial response to hypoxia-ischemic and traumatic brain injuries (each of which can induce dystonia) exacerbates injury and lead to worse outcomes in mouse models ^40,41^. Conversely, inhibition of chronic eIF2α-P pathway activation after the initial response can improve TBI model outcomes ^42^, suggesting impaired regulation of this pathway in either direction may be pathological depending on timing, and could mediate both genetic and environmental causes of dystonia. Therefore, separate from cellular stress, eIF2α-P elevation or depression of could potentially directly alter neuron and circuit properties. Therefore, whether changes in eIF2α-P drive the development of dystonia, play a compensatory role, or represent a nonspecific downstream finding is unknown.

We utilized Drosophila as a genetically tractable model to build on prior work establishing that transgenic *DYT1* flies exhibit dystonia-like movements ^43^. We systematically tested whether altering phosphorylation can directly disrupt normal neuromotor function and which cell types are sensitive to eIF2α-P alterations. We establish both increases and decreases in eIF2α phosphorylation and increased ATF4 as generating dyskinetic movements in flies. We also identify eIF2α-P changes can increase synaptic connectivity, potentially decrease neurotransmission, and pharmaceutical treatment of elevated eIF2α-P with ISRIB may improve some phenotypes.

## Results

### Genetic modification of eIF2α phosphorylation impairs locomotion

We utilized a Drosophila fly model to examine the role of eIF2α-P in locomotion. We started with whole animal partial loss-of-function (LoF) alleles of a major eIF2α kinase (PEK; to increase) and phosphatase (PPP1R15A; to decrease) eIF2α-P (**Fig.1 A**). We confirmed altered eIF2α-P with these genetic manipulations to PPP (**Table 1**) and PEK (**Table 2**) via western blotting (**Supplemental Figure 1**). We identified a decrease in distance traveled in homozygotes for both PPP (**Fig. 1B**) and PEK (**Fig. 1C**). We confirmed that these phenotypes originate from neurons using both the ELAV pan-neuronal and Appl central brain neuron Gal4 drivers for cell-type specific overexpression of both eIF2α-P phosphatase and kinase. We found that neuronal overexpression of either PPP1R15 (PPP) or kinase (PEK) decreased distance traveled (**Fig. 1D-F**). This demonstrates that decreased or increased eIF2α-P levels alone are sufficient to cause locomotor impairments. This is consistent with observations that eIF2α maintains an ideal range of phosphorylation. We reproduced impairments in the locomotor assay with overexpression of the downstream target of eIF2α-P, ATF4, with both ELAV (**Fig. 1G**) and Appl drivers (**Fig. 1H**), suggesting that this phenotype may be attributable, at least in part, to ATF4-mediated gene transcription or repression.

**Fig.1.**
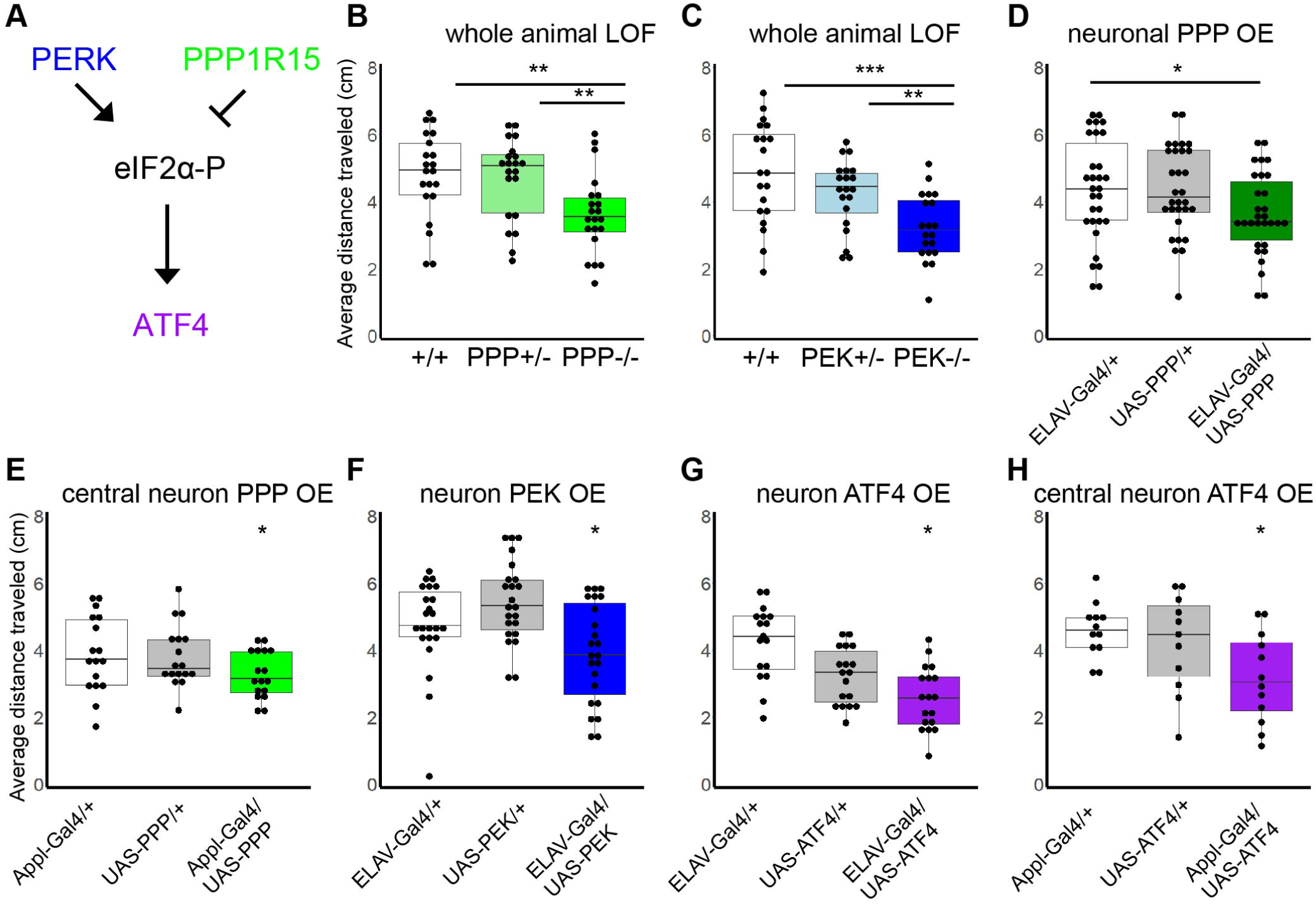
Regulation of eIF2α-P and ATF4 is required for locomotor ability in the fly nervous system. A. Schematic diagram showing regulation of eIF2α phosphorylation (eIF2α-P) by PERK (PEK) kinase and negative regulation by PPP1R15 (PPP) phosphatase leading to expression of ATF4. B-H. Quantification of distance traveled in negative geotaxis assay after 3 seconds. Whole animal hypomorphic loss-of-function variants in PPP (B) and PEK (C) have locomotor impairments. Overexpression (OE) of wild-type PPP1R15 in neurons using an ELAV-Gal4 (D) or central neurons using Appl-Gal4 (E) driver also causes locomotor impairments. F. Neuronal overexpression of wild-type PEK decreases distance traveled. G-H. Overexpression of the downstream mediator of eIF2α-P activity, ATF4, with ELAV (G) and Appl (H) also causes locomotor impairments. Numbers refer to trials where the average distance traveled was calculated for animals of the same genotype, sex, and within the same vial (3-20 flies/vial). Trials are represented by individual dots on the box and whisker graphs. G, n=18. H, n=12. * p<0.05 by paired 2-tailed t-test. For overexpression experiments, statistical significance is represented for both for both genetic controls unless specified with a bar. Additional n values and statistical details in Table 1 (PPP1R15 OE) and 2 (PEK OE).

**Table 1:**
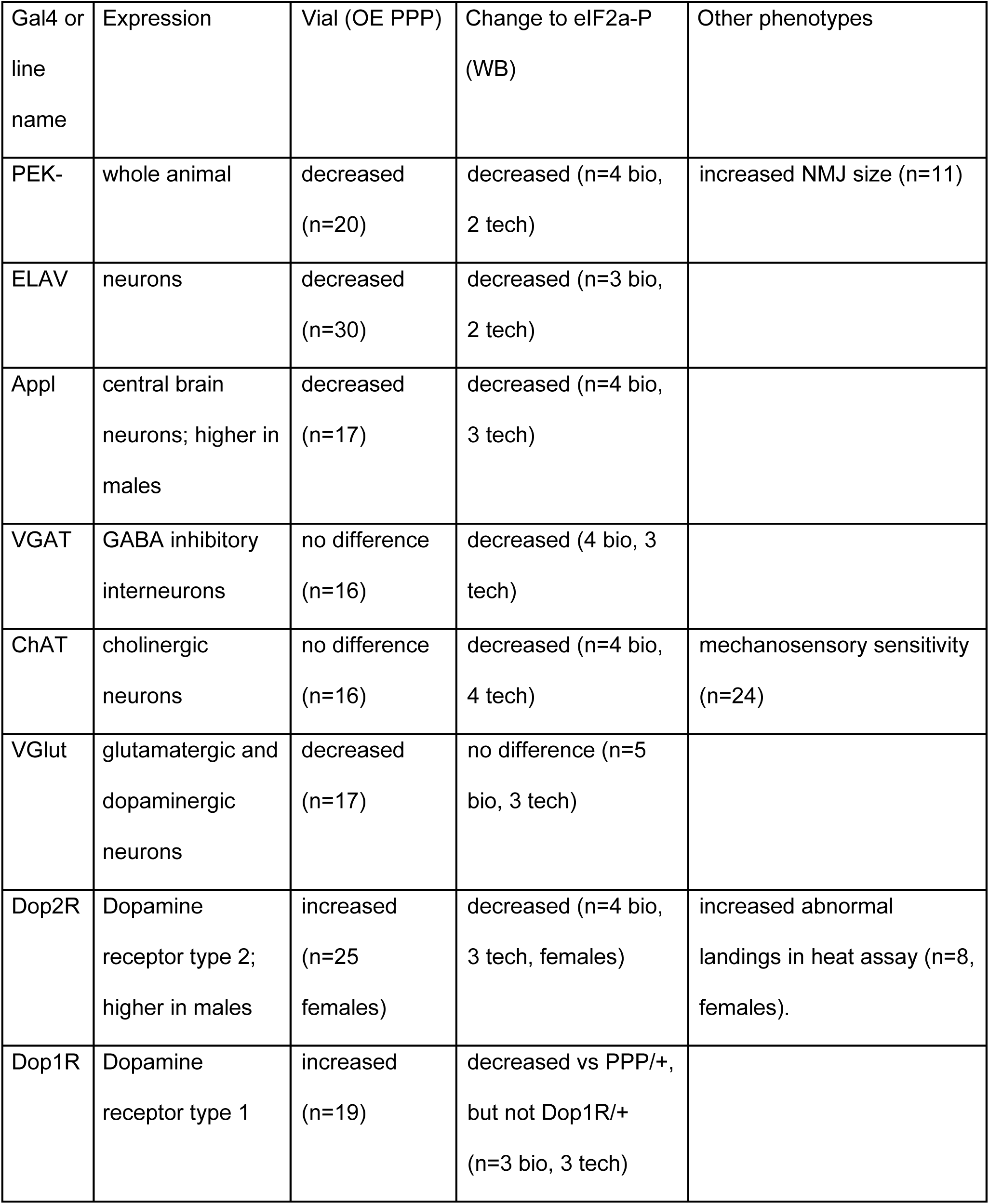
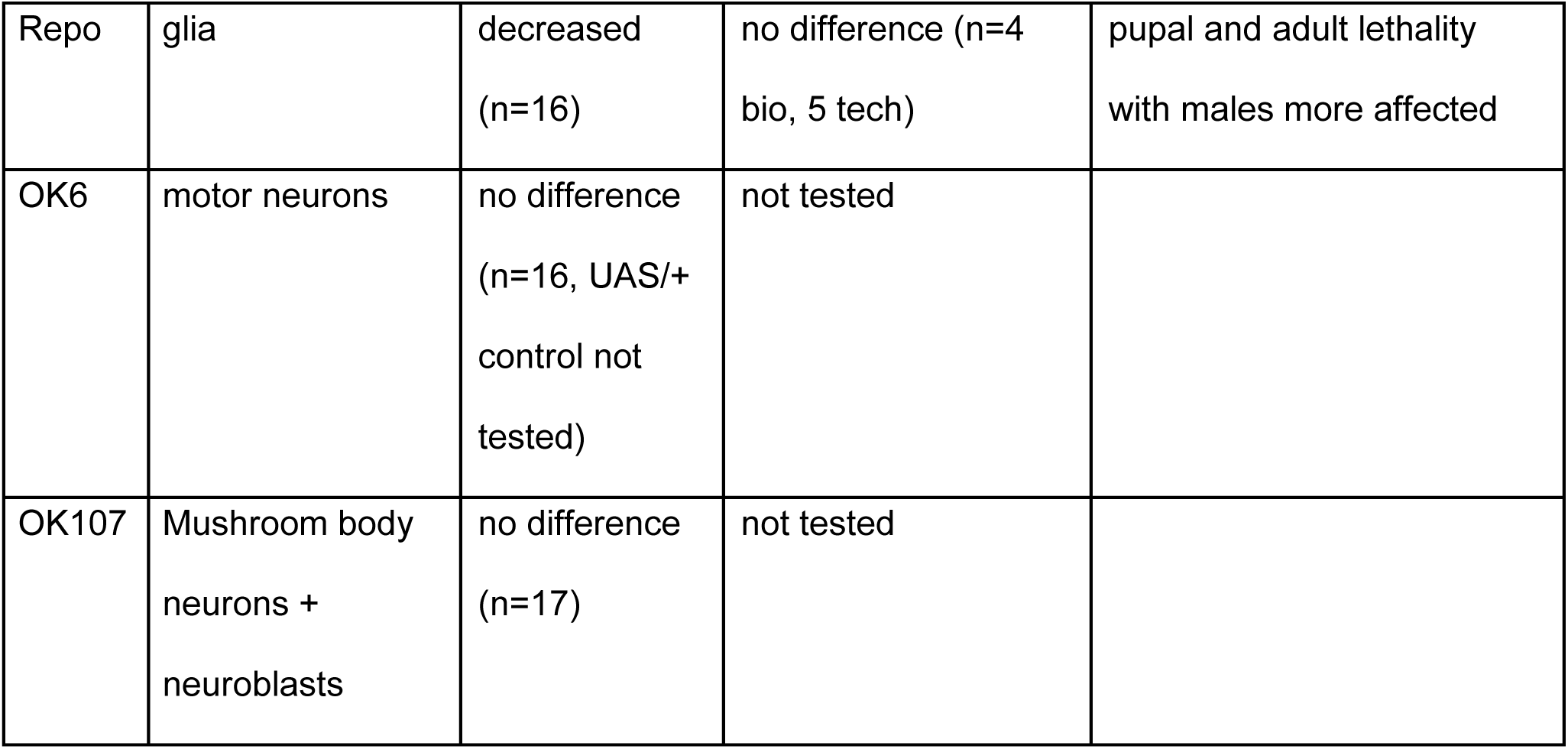
Behavioral and molecular phenotypes from reduced eIF2α-P due to PEK partial LoF or PPP151R overexpression. Bio=independent biological replicates. Tech=samples run on multiple western blots and independently probed. Changes in phenotype required p<0.05 by paired 2-tailed t-test against background genetic control for LoF or both heterozygous genetic controls for overexpression.

**Table 2.**
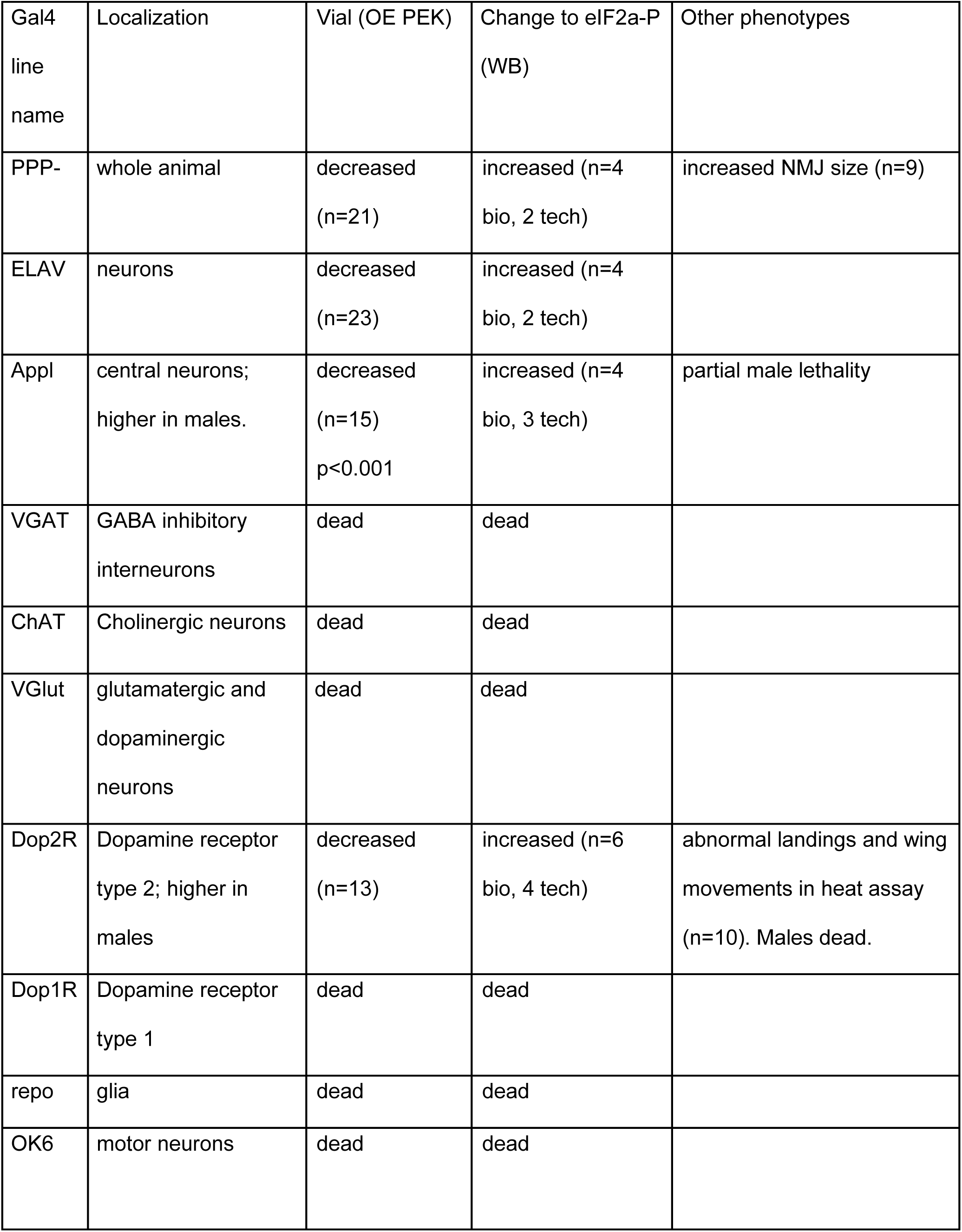
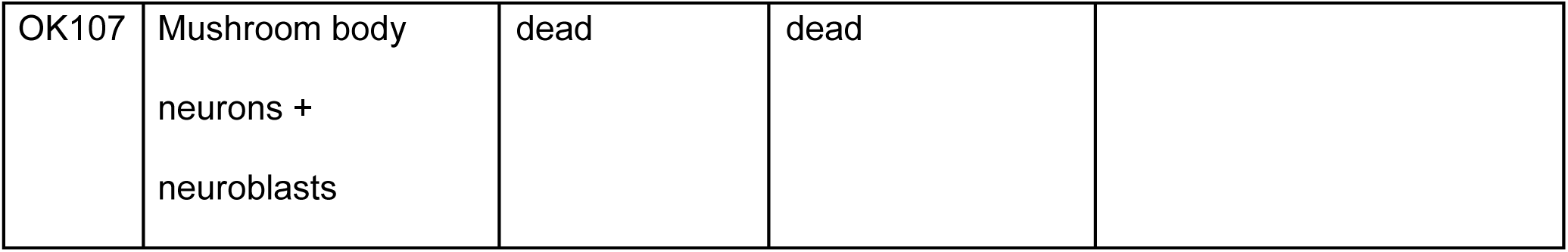
Survival, behavioral and molecular phenotypes from increased eIF2α-P due to PPP151R partial LoF, or PEK overexpression. Bio=independent biological replicates. Tech=samples run on multiple western blots and independently probed. Changes in phenotype required p<0.05 by paired 2-tailed t-test against background genetic control for LoF or both heterozygous genetic controls for overexpression.

| Gal4 line name | Localization | Vial (OE PEK) | Change to eIF2a-P (WB) | Other phenotypes |
| --- | --- | --- | --- | --- |
| PPP- | whole animal | decreased (n=21) | increased (n=4 bio, 2 tech) | increased NMJ size (n=9) |
| ELAV | neurons | decreased (n=23) | increased (n=4 bio, 2 tech) |  |
| Appl | central neurons; higher in males. | decreased (n=15)<br>p<0.001 | increased (n=4 bio, 3 tech) | partial male lethality |
| VGAT | GABA inhibitory interneurons | dead | dead |  |
| ChAT | Cholinergic neurons | dead | dead |  |
| VGlut | glutamatergic and dopaminergic neurons | dead | dead |  |
| Dop2R | Dopamine receptor type 2; higher in males | decreased (n=13) | increased (n=6 bio, 4 tech) | abnormal landings and wing movements in heat assay (n=10). Males dead. |
| Dop1R | Dopamine receptor type 1 | dead | dead |  |
| repo | glia | dead | dead |  |
| OK6 | motor neurons | dead | dead |  |
| OK107 | Mushroom body<br>neurons +<br>neuroblasts | dead | dead |  |

### Locomotor impairments resulting from decreased eIF2α-P can be induced in either neurons or glia, with dopamine-dependent effects

We utilized Gal4 lines with different cell types, neurotransmitter, and region specificity to map the origin of the movement phenotype. Based on findings that the pan-neuronal driver ELAV and Appl impaired locomotion when PPP is overexpressed (**Fig. 1G-H**), we next examined excitatory neurotransmitter subtypes. We again overexpressed the PPP1R15 phosphatase, now in select neuronal subpopulations, and examined alterations to distance traveled (**Fig. 2**), tissue eIF2α-P levels (**Supplemental Table 1**), and survival (**Table 1**). We found decreased distance traveled when expressing PPP1R15 in glutamatergic and dopaminergic neurons (**Fig. 2D**).

**Fig.2.**
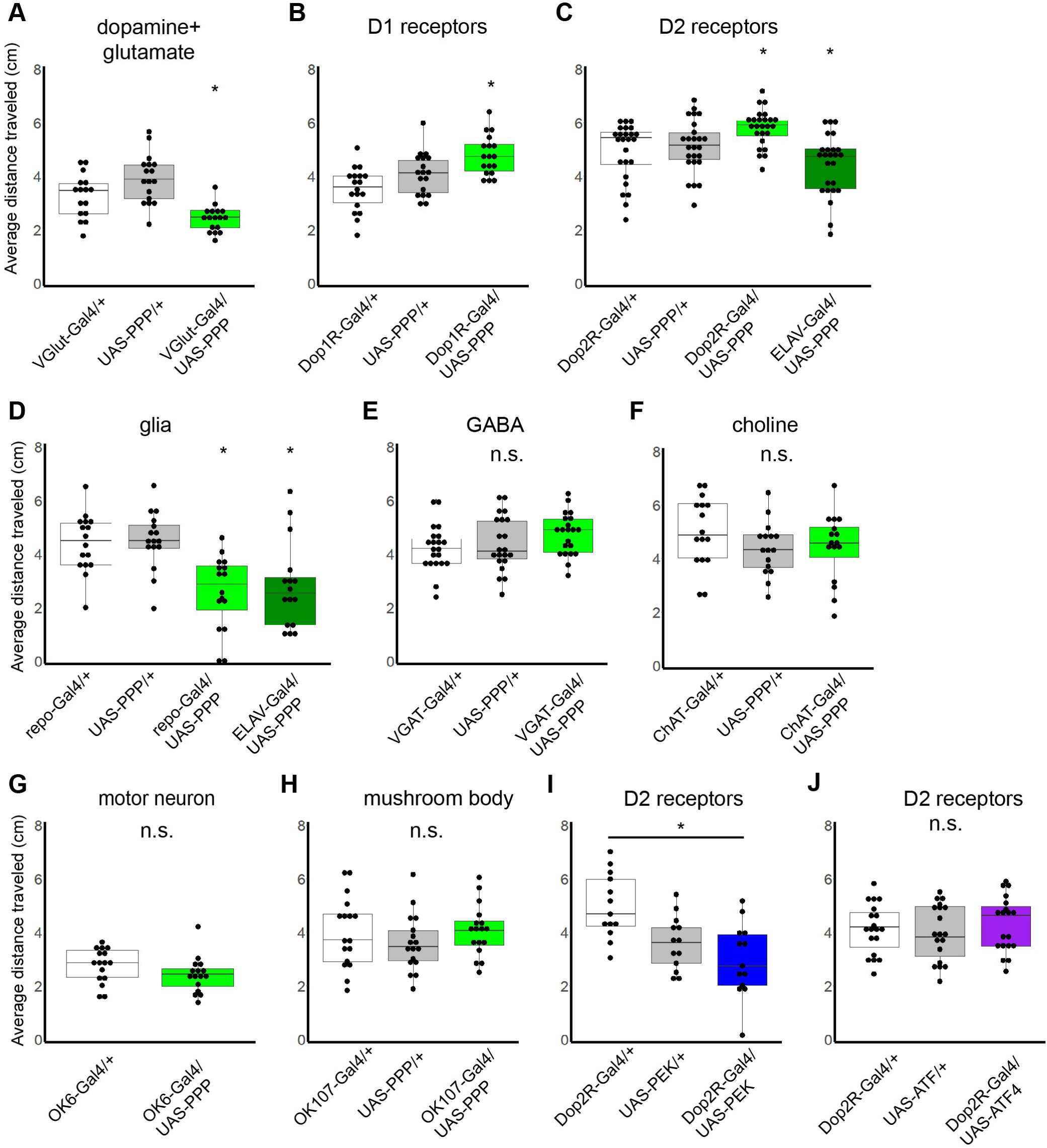
Mapping requirement for eIF2α-P in specific cell types using PPP1R15+, PEK+, and ATF4+ overexpression (OE). A-J. Quantification of distance traveled in flies overexpressing PPP+ (green). Overexpressing PPP1R15 using VGlut-Gal4, which is present in both glutamatergic and dopaminergic neurons, (A) decreases locomotor function. Overexpressing PPP1R15 in neurons expressing dopamine receptors D1 (B) and D2 (C) increased distance traveled. Glial expression (D) was also found to impair locomotor function. There was no change in locomotion with PPP1R15 expression in inhibitory GABA (E), excitatory cholinergic (F), glutamatergic motor (G) or mushroom body lobes, a learning and memory structure in the fly (H). I. Overexpressing PEK+ (blue) in D2-type expressing neurons decreased distance traveled. J. There is no change in distance traveled by overexpressing ATF4+ in D2 neurons (n=19). Numbers refer to number of trials, represented as individual dots on box and whisker plots. * p<0.05 by paired 2-tailed t-test compared to both genetic controls unless specified with a bar. Detailed numbers and statistics in Table 1 (PPP1R15 OE) and Table 2 (PEK OE). n.s. indicates not significant.

We attempted to localize the phenotype to neuronal populations hypothesized to be important for regulating motor function in dystonia. Like humans, flies increase motor activity in response to dopamine with conserved receptors and cellular signaling responses ^44^. We found that the distance traveled increased when we expressed PPP1R15 in both D1-and D2-type dopamine receptor neurons (**Fig. 2B-C).** ^45^. Surprisingly, we also detected a phenotype when expressing PPP1R15 in a pan-glial driver (**Fig. 2D**), indicating that the motor phenotype may not exclusively map to neurons. We found no change in locomotor function when expressed in inhibitory neurons. (**Fig. 2E**) Although there was no change in distance traveled for overexpression of PPP1R15+ in central cholinergic excitatory neuron populations (**Fig. 2F**), we did observe a stronger response to the tap stimulus that warranted follow up. We did not see a locomotor phenotype when PPP1R15 was overexpressed in motor neurons (**Fig. 2G**) or in the mushroom body (**Fig. 2H**), a higher-order brain structure important for initiating and ceasing movements

### Increased eIF2α-P across the nervous system creates developmental lethality and locomotor impairments localizing to D2-expressing neurons

Given that we found that overexpression of PEK-kinase in neurons with the ELAV driver caused locomotor impairments (**Fig. 1F**), we examined the same battery of Gal4 lines and found overexpression of PEK in most cell types caused early developmental lethality (**Table 2**). The requirement for PPP1R15 and PEK during development has been described previously ^9,46^, but this demonstrates sensitivity to overexpression of PEK in multiple neuronal subpopulations. Interestingly, female Dop2R-PEK+ progeny survived and had decreased distance traveled in a locomotor assay (**Fig. 2I**) compared to increased distance in Dop2R-PPP+ (**Fig. 2C**). However, Dop2R overexpression of ATF4 did not alter distance traveled (**Fig. 2J**). This is the only bidirectional phenotype observed in these studies, demonstrating a unique relationship between the levels of eIF2α-P in D2-type neurons and motor control.

The Dop2R-Gal4 and Appl-Gal4 insertions are located on the female sex chromosome and are expressed twice as much in males compared to females. These 2 lines had female progeny surviving to adulthood, but males failed to survive. Therefore, PEK overexpression phenotypes are dose-dependent and not due to dominant effects of the P-element insertion.

### Neurons expressing D2-receptors have impaired motor coordination evoked by heat with increased and decreased eIF2α-P, ATF4 overexpression, and the dystonia risk allele htorΔE

Human dystonia can be precipitated by sensory stimuli such as heat ^47^, mechanosensory sensation, positive or negative emotion, or passive or volitional movement ^48^. Dystonia-like movements manifest in flies with mutations in orthologs of dystonia genes *ATP1A3* and *DYT1* as hypokinetic or hyperkinetic repetitive movements evoked by mechanical and/or thermal stimulation ^43,49^. At elevated temperatures below heat stress, this increase in temperature is thought to increase synaptic transmission and excitatory neuronal drive due to flies being a poikilothermic model ^50^. Dopamine deficient states can lead to dystonia; tyrosine hydroxylase and GTP cyclohydrolase deficiency can lead to dystonia and treatment with L-Dopa can reduce dystonia ^25^. We thus examined D2-expressing neurons, which is where bidirectional sensitivity to eIF2α-P levels were observed, and which are important for suppressing movements as part of the indirect pathway ^19^. We overexpressed PEK to see if modifying eIF2α-P in D2-expressing neurons can manifest features of dystonia, such as dyskinesia or disruptions to volitional movement.

We captured videos over 3 minutes at 37-40°C, a physiological temperature below the threshold to induce heat shock or paralysis from action potential propagation failure ^51^. We observed hyperkinetic movements in flies subjected to elevated temperatures that manifested as wing flapping in absence of flight that were not present in either genetic control. Events where animals that had wings in motion for >4 video frames while standing or walking were quantified. 82% (290/352) of bouts were 1 second or shorter. These bouts were present in animals expressing PEK+ in D2-receptor cells at a rate of 7.6 events/fly on average (**Fig. 3A**). Since these wing movements were not present in controls, we conclude that this was not an attempt at cooling. The elevated temperature also increases the number of fly movements, measured as flights, jumps, or drops. The frequency increases and then plateaus over time, consistent with heat-driven increase in activity followed by acclimation (**Fig. 3B**). The number of movements increased equally in D2-PEK overexpressing flies and their genetic controls (**Fig. 3C**). However, Drosophila overexpressing PEK in D2 receptor neurons had difficulties in coordinating landings, defined as instances wherein an animal landed on its back, side, or abdomen (**Fig. 3D**), with most abnormal landings involving contact between the side or back with the floor. We observed a 3-fold increase in percent of landings which were abnormal in genotype-blinded scoring in flies with D2-neuronal elevated eIF2α-P compared to their genetic controls per total attempted landings (**Fig. 3E**). Often, flies landing on their back or side had difficulty righting as well and required several attempts to return to standing (**Supplemental Video 1**). We identified good agreement between independent observers (Gwet AC1 95% CI 0.87-0.91, p<10^-16^) in identifying these abnormal landings. D2 overexpression of the eIF2α-P downstream effector ATF4 also increased the percentage of abnormal landings (**Fig. 3E**). Surprisingly, decreasing eIF2α-P by overexpression of eIF2α phosphatase, PPP+ also increased the percentage of abnormal landings (**Fig. 3F**). This suggests that for this phenotype, altered phosphorylation in either direction is pathological.

**Fig 3:**
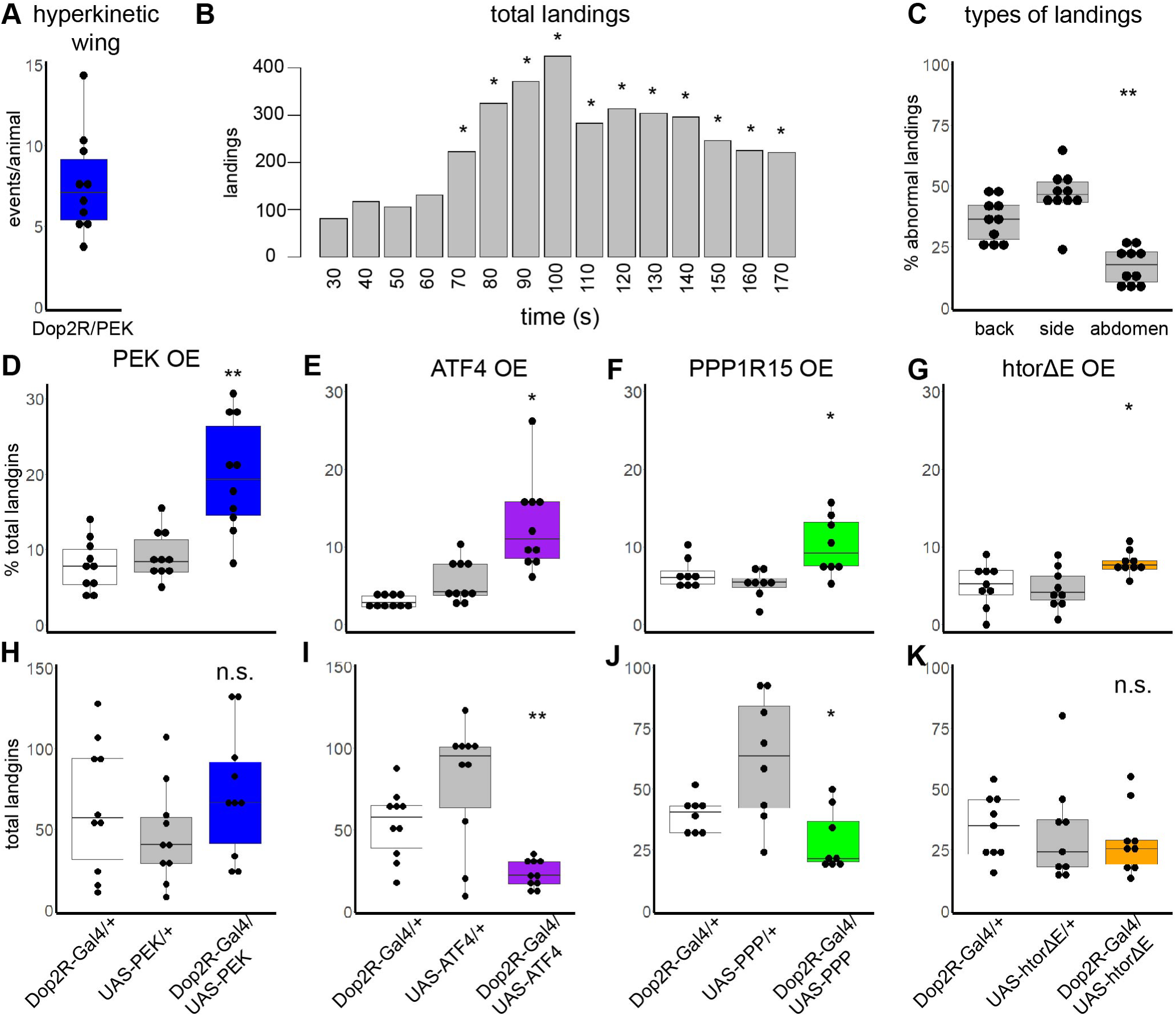
Heat-evoked movements due to increased and decreased eIF2α-P, ATF4 OE, and htorΔE in D2 receptor expressing neurons. A. Hyperkinetic wing events per animal, defined as flapping for >4 video frames in the absence of flight, were only observed in Dop2R/PEK flies. B. The number of flights and jumps increases significantly with exposure to 37-40°C temperature in Dop2R/PEK compared to activity at t=30 seconds. C. Distribution of the body part of the flies involved in abnormal landings. The back and side of the animal contacted the floor significantly more often than the abdomen. D-G. Percent of landings scored as abnormal due to contact between floor and the fly’s abdomen, side, or back was increased in D2-receptor overexpression of PEK (D), ATF4 (E), PPP1R15 (F), and htorΔE (G). H-K. Total landing attempts. There is no difference in total landings between D2 PEK OE and genetic controls (H). Total landing attempts were decreased in D2 OE of ATF4 (I) and PPP (J), indicating reduced flights and activity. There was no difference in total landing attempts in htorΔE (K). Notably, OE did not increase the number of attempted flights in any case, indicating coordination defects are not just due to hyperactivity. A, D, H n=10 trials, females only. ** p< 0.001. B. n=10 trials, 3663 total landings, D2 PEK OE only. * p<0.05 2-tailed t-test vs at baseline measured between 30-40 seconds after starting video. C. n=10 trials, 628 total abnormal landings, D2-PEK only. ** p=0.001 abdomen vs. back and p=0.0002 abdomen vs. side calculated by 2-tailed t-test. E, I. n=10 trials, female only. * p < 0.002, ** p<0.005. F, J. n= 8 trials female only. * p<0.03. G, K. n=9 trials, males only. * p<0.04. Statistics calculated using paired 2-tailed t-test requiring p<0.05 compared to both genetic heterozygous controls. n.s. indicates not significant. An average of 9 flies/trial were used.

We hypothesized that similar dyskinetic movements may be present in a model of primary genetic dystonia when expressed in the same cell type and subjected to the same assay. We next examined a fly ortholog of the *DYT1* risk allele, which features a deletion of glutamic acid at position 302/303 (htorΔE). Flies expressing htorΔE in neurons, dopaminergic neurons, and muscles are paralyzed after extended exposure to very high temperatures^43^. We observed D2 localized overexpression of htorΔE increased the percentage of abnormal landings with no evidence of paralysis (**Fig. 3G**), suggesting abnormal landings are likely a coordination defect rather than due to an aspect of the paralysis phenotypes previously observed. No abnormalities were noted when overexpressing wildtype *DYT1* (htorA; data not shown).

Some genotypes appeared to have fewer attempted flights, so we also quantified the total landings per fly. Total landings were not altered with D2 localized overexpression of PEK (**Fig. 3I**), while the landings per fly decreased for ATF4 (**Fig. 3J**), decreased for PPP (**Fig. 3K**), and did not change for htorΔE (**Fig. 3L**). Therefore, overexpression of ATF4 and PPP decreases the tendency of the flies to initiate flight. Hyperkinetic wing movements were only detected in the Appl-PEK genotype and was not present when overexpressing ATF4, PPP, or htorΔE. We also did not observe an increase in non-evoked coordination issues, such as falls or alterations in stance. This suggests the increased proportion and non-purposeful quality of the movements we observed with overexpression of PEK, PPP, ATF4, and htorΔE in D2-receptor neurons were truly dyskinetic (abnormal, unwanted movements) and did not simply represent generalized hyperactivity or motor impairment.

Thus, flies with elevated eIF2α-P and ATF4 manifest abnormal control of complex motor program required for successful landings; excessive eIF2α-P is additionally associated with hyperkinetic wing movements.

### Reduced eIF2α-P, overexpression of the htorΔE dystonia risk allele and ATF4, or decreased neurotransmission in cholinergic neurons increases hyperkinetic responses to mechanosensory stimulation

Mechanical stimulation evokes hyperkinetic movements in Drosophila with mutations in dystonia-associated genes ^49^ known to increase inhibitory neuron excitability and firing ^52^. We examined overexpression of the eIF2α phosphatase, PPP1R15+, in cholinergic neurons because of the observed sensitivity to tapping in the locomotor assay. Additionally, striatal acetylcholine appears to play an important role in dystonia ^25^ with increased cholinergic activity in *DYT1* animal models ^27^ and clinical response to anticholinergics. To determine sensitivity to a mechanosensory stimulus, flies were vortexed for 10 seconds and time to recover was recorded. We found that ChAT/PPP+ flies had increased duration of hyperkinetic movements after mechanical stimulation, including wing flapping without flight and rapid leg movements from a prone position, as well as unproductive movements, such as repeated attempts to stand resulting in falling and further hyperkinetic movements (**Fig. 4A; Supplemental Video 2**). Although a similar assay has been used for studying epilepsy, these movements appear distinct from a seizure as they are not periodic or oscillatory and do not have a refractory period (not shown). Although we were not able to test overexpression of PEK+ due to lethality (**Table 2**), we overexpressed ATF4, the downstream mediator of elevated eIF2α-P, and observed increased time to recover (**Fig. 4B**).

**Fig.4.**
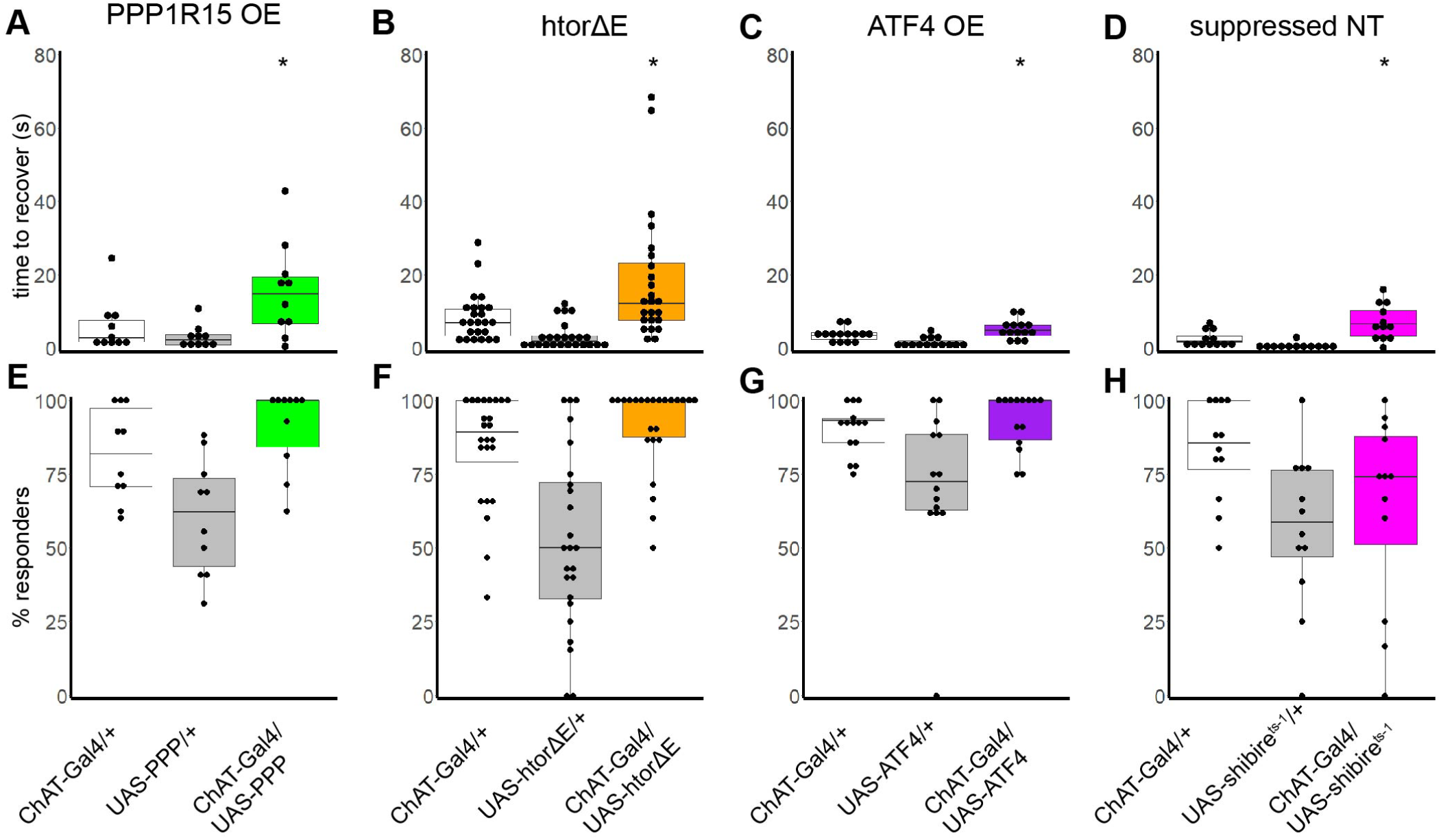
Sensitivity to mechanical stimulation with cholinergic expression of PPP1R15+, htorΔE, ATF4, and suppressed neurotransmission (NT). A-D, Quantification of recovery time. Expression of PPP1R15+ (A) and htorΔE DYT1 dystonia risk allele (B) in cholinergic neurons show increased time to recover from a mechanosensory event compared to controls. Expressing a wildtype DYT1 allele, htorA does not change recovery compared to controls (**Supplemental Figure X**). E. Overexpression of wild-type ATF4 also increases time to recovery. D. Temperature sensitive dynamin (*shibire^ts-1^*) decreases dynamin-mediated endocytosis, synaptic vesicle recycling, and ultimately neurotransmission. Decreasing neurotransmission in cholinergic neurons also increases the time to recover from mechanical stimulation (D). E-H, Quantification of percent of animals responsive to mechanosensory stimulus, defined flies exhibiting hyperkinetic movements at the bottom of the vial. Non-responders were counted as flies walking on the vial wall when the vials entered the camera field of view. There is no difference in the proportion of flies responding to the mechanical stimulus when compared to both heterozygous controls. * p≤ 0.05, paired 2-tailed t-test. Numbers indicate trials, represented as individual points on the box and whisker plot. An average of 9 flies/trial were used.

We asked whether overexpression of the htorΔE dystonia risk allele in cholinergic neurons similarly increased sensitivity to a mechanical stimulus. Like flies overexpressing PPP1R15+, htorΔE flies had hyperkinetic and unsuccessful movements with mechanical stimulation, with longer time to recover (**Fig. 4C**). In comparison, overexpressing wildtype human torsin A (htorA) as a control did not alter time to recover compared to control animals (**Supplemental Figure 2**).

We hypothesized that reductions in eIF2α-P may be altering neuronal or circuit properties such as neurotransmission. Cell-intrinsic expression of htorΔE has been demonstrated to reduce dopamine neurotransmitter levels in a mouse model ^53^. To determine if the mechanosensitivity phenotype could be due to altered neurotransmission, we expressed a temperature sensitive dynamin (*shibire^ts-1^*) in cholinergic neurons. This temperature sensitive mutation blocks endocytotic recycling and subsequently reduces neurotransmission at elevated temperature. Flies were reared at low temperature (18°C) and moved to room temperature for recording (22-23°C) but kept below the restrictive temperature which induces paralysis (30°C) ^54^. Animals reared this way were healthy, with no observed motor defects at room temperature and no developmental or early adult lethality (not shown). In response to mechanical stimulus, flies expressing shibire with predicted decreases in cholinergic neurotransmission exhibited the same hyperkinetic movements as overexpressed PPP1R1R+ and htorΔE. In addition to a longer time to recover (**Fig. 4D**), we observed infrequent, temporary paralysis thought to be due to increased muscle activation because paralysis was both preceded and followed by hyperkinetic movements. The proportion of animals exhibiting dyskinetic movements in response to the stimulus did not change from ChAT/+ controls in any experiment (**Fig. 4E-H**), indicating the difference was due to stronger and longer lasting hyperkinetic responses rather than a change in the number of flies which responded. These results show that decreased eIF2α phosphorylation, ATF4 and htorΔE expression, and decreased neurotransmission in cholinergic neurons increase sensitivity to mechanically elicited dystonia-like movements.

### Increased or decreased eIF2α-P increases axon terminal size and bouton number at the neuromuscular junction with dysregulated synaptic structure

Since eIF2α-P modifications induce heat-(**Fig. 3**) and mechanosensory-evoked (**Fig. 4**) dyskinetic movements and increased eIF2α-P has been observed to decrease dendrite arbor size ^55^, we asked if eIF2α-P levels regulate neuron connectivity. Increased connectivity between brain regions have been identified in functional imaging dystonia patients ^17,18^. Further, alterations to synaptic plasticity are observed in dystonia models including a decreased threshold for LTP and absent LTD in a *DYT1*/htorΔE mouse ^22^ and impaired LTD with intact LTP in *DYT6* mouse ^23^. Reduced dopamine release has been observed in ATP1A3 shRNA knockdown mouse model ^52^ suggesting that synaptic release and connectivity may be altered in dystonia. Finally, an increase in the number of synaptic release sites was identified in Drosophila motor neuron terminals at the neuromuscular junction (NMJ) with both overexpression and knockdown of the fly ortholog of *DYT1, dtor* ^56^. To determine if alterations in synaptic connectivity occurs with genetic manipulation of eIF2α-P levels, we examined the NMJ using immunostaining (**Fig. 5A-B**). We assessed a whole-animal, partial LoF using hypomorphic alleles of both *PPP1R15* and *PEK*. We found increased axon terminal size relative to muscle area in both PPP1R15^-/-^ and PEK^-/-^ homozygotes compared to genetic controls (**Fig. 5C**). Close examination of the boutons, the specialized structure where synaptic release occurs, revealed irregular bouton structure and density (**Fig. 5F-G**), with the appearance of satellite boutons budding from the main bouton. Bouton number was significantly increased in both PPP1R15^-/-^ and PEK^-/-^ homozygotes (**Fig. 5H**). Therefore, increases in synaptic connectivity may arise from alterations to eIF2α phosphorylation and could potentially underlie changes to motor control.

**Fig.5.**
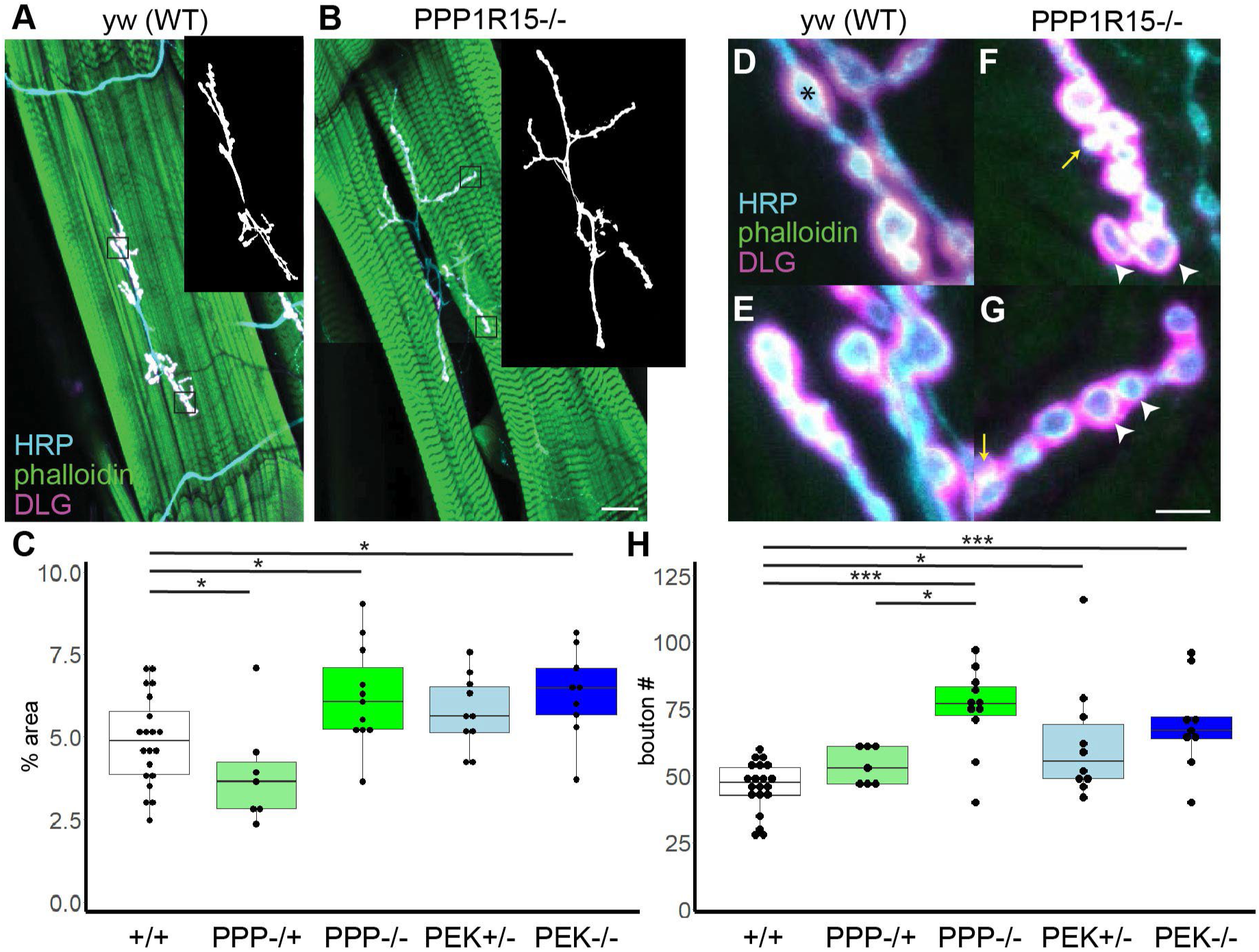
Altered eIF2α-P by reductions in PPP1R15 or PEK increases axon terminal size at the neuromuscular junction (NMJ). A-B Sample images reconstructed from confocal stacks from third instar larva A3 6/7 NMJs. Muscles visualized by phalloidin binding filamentous actin in green, neuron membranes recognized by HRP in cyan, and postsynaptic density recognized by DLG in magenta. Inset, neuron membrane visualized by anti-HRP post-threshold processing during quantification with ImageJ. Boxes, regions magnified in D-G. A, *yw* genetic control. B, *PPP1R15^G18907^* loss-of-function hypomorph homozygote (-/-). Scalebar=50 μM. C, quantification of the neuron area (HRP) normalized to 100 μm^2^ muscle area. D-G, High magnification images of synaptic boutons, with characteristic rounded synaptic release site (cyan) surrounded by postsynaptic density (magenta). D-E wild type boutons (asterisk) are relatively uniform in size and arranged evenly along the axon length. F-G *PPP1R15^G18907^* boutons are densely packed (white arrowheads) and feature satellite boutons budding off main boutons (yellow arrows). Scalebar=10μm. H, Quantification of bouton number. Numbers indicate the number of NMJs. +/+ indicates homozygous yw or w^1118^ controls (n=20). PPP-/+ heterozygotes n=7, PPP-/-homozygote n=11, PEK-/+ heterozygote n=9, PEK-/-n=9. HRP=horseradish peroxidase. DLG=discs large, psd95 ortholog. *p<0.02 and ***p<0.0002 by Mann-Whitney rank sum test.

### Pharmacologic treatment of eIF2α-P leads to partial phenotypic rescue in vivo

We investigated whether bypassing the translational suppression caused by elevated eIF2α-P could improve outcomes in the fly model. We first determined that there was no change in distance traveled in genetic controls treated with vehicle or 500 nM ISRIB (**Fig. 6A**). We overexpressed PEK in central brain neurons with the Appl driver and treated animals with either ISRIB or vehicle control during larval development. We also did not detect a difference in distance traveled between vehicle-treated and ISRIB-treated Appl-PEK+ expressing animals (**Fig. 6B**). We next established that 500 nm ISRIB did not create developmental (not shown) or early adult lethality (**Fig. 6C**) in genetic controls. However, treatment with ISRIB during development significantly improved adult lifespan of flies with neuronal overexpression, Appl-PEK+ (**Fig. 6D**). This suggests that pharmaceutical modification of elevated eIF2α-P may improve some phenotypes but may be sensitive to timing window or work best within certain ranges of phosphorylation ^57^. Notably, treatment with ISRIB did not cause any observable phenotypes for control flies.

**Fig. 6.**
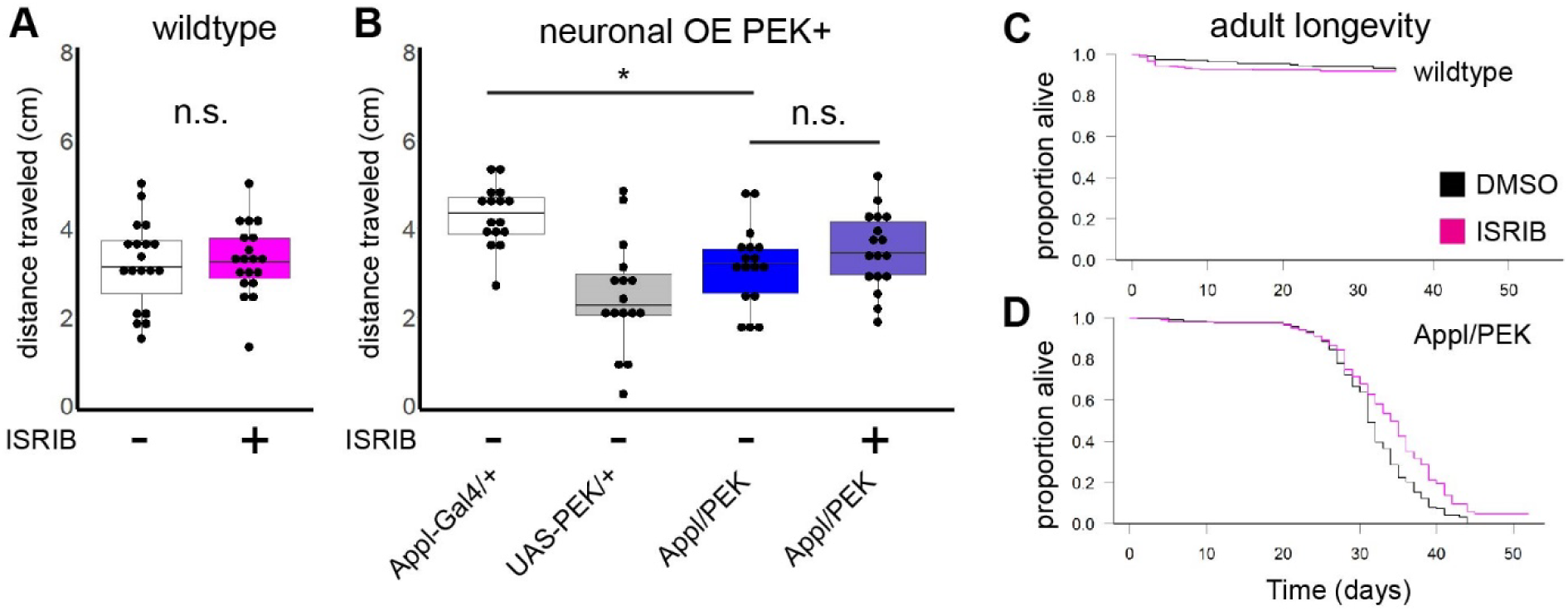
Pharmacological treatment with ISRIB partially rescues phenotypes of elevated eIF2α-P. A-B, Average distance traveled in adult locomotor assay. Treatment with 500 nM ISRIB during development did not change average distance traveled in genetic controls (A) or Appl-PEK+ overexpressing (OE) animals (B). C-D, Adult survival. There was no change in survival in wildtype genetic control flies when treated with 500 nM ISRIB during development (C), however, treatment significantly improved adult lifespan in animals with neuronal overexpression of PEK+ and elevated eIF2α-P (D). * p<0.003 2-tailed t-test. Survival difference p= 7e-05 calculated. n.s. indicates not significant.

## Discussion

The regulation of the eIF2α phosphorylation is tightly controlled and is emerging as a candidate mechanism for dystonia of diverse origins ^31^. In these studies, we investigated whether eIF2α-P signaling would be sufficient to explain motor features of dystonia using a Drosophila model. Consistent with this, we found eIF2α-P is required for development of normal motor function in both glia and neurons, with further localization to cholinergic, dopaminergic/glutamatergic, and dopamine-receptor expressing neurons. Altered eIF2α-P signaling in D2-receptor expressing neurons and cholinergic neurons increases susceptibility to hyperkinetic and unproductive movements in the presence of heat and mechanical stimulation, respectively. We identified overlap with both phenotypes using an independent dystonia model, the ΔE allele of torsin/DYT1 ^43,58^. Mechanosensitivity was also achieved with suppressed neurotransmission using a dynamin mutant; interestingly, variants in dynamin have also been identified as a genetic cause of dystonia ^59^.

We identified increased synaptic terminal size in both PPP1R15 and PEK partial LoF variants that increased and decreased eIF2α-P, respectively. We also identified increased bouton density and satellite boutons. Notably, both LoF and OE of Drosophila torsin creates a similar increase in the number of boutons ^56^ and satellite boutons are associated with mutations in endocytosis, including shibire/dynamin ^60^. These findings demonstrate that bidirectional changes to eIF2α-P, dysregulated DYT1, and suppressed endocytotic recycling driving the availability of synaptic vesicles are phenotypically similar at both the synaptic and movement levels. Together, these results suggest either increased or decreased eIF2α-P can may create cell-type specific alterations to connectivity or neurotransmission relevant to dystonia.

Dysregulated eIF2α-P within D2-expressing neurons fails to suppress abnormal movements when excitatory drive is high. This is consistent with findings that D2-antagonist compounds can elicit tardive dyskinesias ^25^. Activation of D2 receptors expressed in the indirect pathway decreasing activity of these neurons which inhibit motor activity, and the loss of this suppression has been identified in various hyperkinetic movement disorders ^25^. Activation of D1 receptors while suppressing D2 receptors subsequently increases motor drive in focal dystonias ^61^. Consistent with our finding that both D1 and D2 receptors are sensitive to eIF2α-P perturbations, overexpressing PEK specifically in dopaminergic neurons reduces distance traveled in a locomotor assay in older Drosophila and those subjected to chronic heat stress ^62^. Knocking down PEK in mouse dopaminergic neurons also increase motor activity ^63^. Together this demonstrates that eIF2α-P dysregulation in both dopamine releasing neurons and neurons modulated by dopamine, especially those expressing D2 receptors, can lead to alterations in motor control.

We also found increased sensitivity to mechanical stress, but only in the cholinergic neurons. Conversely, we observed heat did not appear to affect coordination of landing when altering eIF2α-P or htorΔE in cholinergic neurons. We were also unable to examine the hypothesis of reduced neurotransmission in the heat assay due to the temperature sensitive nature of the genetic reagent. In vertebrates, alterations to acetylcholine in the striatum regulating dopamine release is a candidate for dystonia pathophysiology ^27^. In insects, acetylcholine is a central excitatory neurotransmitter with glutamate neurotransmission at the NMJ ^64^. In both vertebrates and insects, acetylcholine plays an important role in inhibitory control for suppressing movements ^65^. One possible mechanism is acetylcholine activation of mAChRs facilitates dopamine release in the striatum ^66^ which inhibits neurons expressing D2 receptors. D2-expressing neurons subsequently play a key role in motor inhibition, with abnormal D2 signaling contributing to both tardive dyskinesias and dystonia ^25^. This suggests a model where decreased neurotransmission in cholinergic neurons or D2-receptor expressing neurons can uncover dystonia-like hyperkinetic movements or other motor impairments.

Studies of the motor circuit in dystonic patients have identified altered connectivity between sensory-motor cortex, thalamus, cerebellum, and basal ganglia ^17,18^. It is hypothesized that disrupted synaptic plasticity, connectivity, neurotransmission, or neuromodulation may underlie the circuit changes. Heat and mechanical stimulation in Drosophila increase global neuronal activity and likely represent increased excitatory inputs onto sensitized cell types, resulting in hyperkinetic motor activity. We also observed increased synaptic connectivity with alterations to eIF2α-P. Therefore, the molecular changes from increased/decreased eIF2α-P and htorΔE could contribute to both reduced neurotransmitter release within the indirect pathway for inhibiting unwanted movements and abnormally increased synaptic connectivity elsewhere. Consistent with this, both decreased calcium transients ^33^ and enhanced LTP ^22^ have been observed in *DYT1* models.

We identified the overexpression of ATF4, a transcription factor translated with elevated eIF2α-P signaling, was also sufficient to generate motor impairments in locomotor, heat, and mechanical stress assays. ER stress from misfolded proteins ^67^, regulators of eIF2α-P such as PRKRA ^37^, and downstream factions including ATF4 ^32^ are being identified as individual genetic causes of dystonia. Future studies could examine which ATF4 targets are abnormally regulated in dystonia and whether they would be a potential target for treatments. Modification of eIF2α-P with compounds such as ISRIB may hold promise in the future for treating dystonia, however, our findings suggest that timing windows and levels of eIF2α-P would be important considerations for an effective treatment.

## Acknowledgments

We are grateful to Nahla Haque, Megan Kotzin, James Liu, Rosario Padilla Sánchez, Andrew Musmacker, Frances Nowlen, Tristan Shackelford, and Bryce Wilson for assistance with Drosophila rearing, assays, and western blotting. We thank Daniela Zarnescu and Robert Kraft for their advice and training.

We are grateful to Naoto Ito for the kind gift of htor fly stocks and FlyORF for ATF+ and PPP+. The remaining stocks were obtained from the Bloomington Drosophila Stock Center (NIH P40OD018537). The monoclonal antibody developed by C. Goodman was obtained from the Developmental Studies Hybridoma Bank, created by the NICHD of the NIH and maintained at The University of Iowa, Department of Biology, Iowa City, IA 52242. Biomedical Imaging Core microscopy facilities at the University of Arizona College of Medicine – Phoenix were used in these studies.

SL is supported by the CPARF PRG06621 and MCK is supported by NIH 1R01NS106298 and 1R01NS127108.

## Declaration of Competing Interests

The authors declare no competing interests. The funding agencies did not play a role in experiment design or preparation of this manuscript.

## Author contributions

Conceptualization, S.A.L., M.C.K; Methodology, S.A.L.; Investigation, S.A.L., J.F., J.T., R.S., Z.T.; Writing—original, S.A.L; Writing-Review & Editing, all; Funding Acquisition, S.A.L., M.C.K.

## Methods

### Drosophila genetics and rearing

Drosophila reared on a standard cornmeal, yeast, sucrose food from the BIO5 media facility, University of Arizona. Stocks for experiments were reared at 25°C, 60-80% relative humidity with a 12:12 light/dark cycle. Cultures for controls and mutants were maintained with the same growth conditions, especially the density of animals within the vial. The GAL4-UAS system was used to drive cell-type specific expression of eIF2α-P genetic modulators ^68^. For experiments using whole-animal partial loss-of-function alleles, controls were *yw* homozygotes and heterozygotes from the cross between *yw* and the loss-of-function homozygous stock. For Gal4-UAS experiments, controls were heterozygotes from the Gal4 and UAS lines crossed with w^11181^.

A complete list of genotypes, source, and expression information can be found in **Supplemental Table 1**. *w^1118^* was outcrossed with Canton-S and backcrossed with *w^1118^* for 12 generations while selecting for red eyes. UAS-crc (ATF4+) was acquired from FlyORF ^69^. The creation of htorΔE and htorA were described in ^58^ and were a kind gift from Naoto Ito. The remaining lines were acquired from Bloomington Stock Center (NIH P0OD018537).

### Western blotting and quantification

Protein samples prepared from heads of 10-20 males and females in protein extraction buffer plus 1% protease inhibitor (ThermoFisher) and 1% phosphatase inhibitor (Sigma) as previously described ^70^. Western blotting performed according to standard methods with detection on a 0.2 µm nitrocellulose membrane. For eIF2α, phosphorylation antibody was imaged first, then the membrane was washed and reprobed for total eIF2α. Antibodies used for study are listed in **Supplemental Table 2**. eIF2α-P: 1:1000 rabbit α phospho-S51 (Cell Signaling 3597), 1:500 rabbit α eIF2S1 (Abcam 4837) and detected with a 1:10000 goat anti rabbit secondary antibody conjugated with HRP and imaged with ECL (GE healthcare NA931, NA934).

Chemoluminescence captured using FluorChem Imager (Biotechne) and quantified with Image studio Lite (LiCor). eIF2α phosphorylated/total ratio was normalized to Gal4/+ heterozygous control for each biological replicate. The same size area was used to measure total protein and puromycin signal for each lane to reduce variability. At least 3 technical replicates and 4 independent biological samples were used for quantification. Differences between genotypes calculated using 2-tailed paired t-test statistic and graphs generated using R (4.2.2). Software packages used to analyze data can be found in **Supplemental Table 3**.

### Drosophila movement assays

We used naïve, unmated flies collected as pharate adults and sequestered for 14 days before recording. Flies were adapted to room conditions for 1 hour before moving to empty assay tubes or petri dishes in groups of 10-20. All assays included sex-matched genetic controls as part of the trial. Flies of both sexes were used for all experiments except for Dop2R-PEK and Appl-PEK due to male lethality. Video recorded with Canon EOS Rebel T7 camera at 60 fps and visualized with Adobe Premier Pro or Rush.

Distance traveled assay was performed using paired, coded vials of control and mutant flies ^71^. Distance measured from still image from video at 3 seconds post-tapping using ImageJ (NIH, v.1.50i) to measure distance function from middle of fly thorax to bottom of vial. Scorers were blind to genotype.

The adult heat assay was adapted from ^43^ with scoring of the video to quantify hyperkinetic and uncoordinated movements. Groups of flies were loaded via aspirator into 3 cm petri dishes, matched for number of flies and sex for each recording. Damp filter paper was used to regulate humidity in heat assay chamber. Temperature was regulated by water bath with 37-40°C temperature continuously monitored. Video recorded for 3 minutes after assay chamber submersion. Hyperkinetic wing events were defined as wings in motion for >4 video frames during a non-flight behavior such as walking or standing; flies landing or initiating flight afterwards were excluded. Videos were scored for the number of total floor landings (defined as any action where a fly contacted the floor, such as after a drop, flight, or jump from any surface of the petri dish. Abnormal landings were defined as when the fly contacted the floor with the caudal tip of the abdomen, side, or back. Abdomen landings required the fly to be perpendicular to the floor with no legs in contact. Side landings required the fly to have all six legs visible and more than 2 video frames to reach an upright position. Two independent scorers were used to develop criteria for scoring and for scoring Dop2R-PEK. Multiple iterations of training, feedback, and criteria clarification were used until >90% agreement was achieved for all 10 trials. Scorers were blind to genotype during scoring total and abnormal landings. The percentage of abnormal landings was calculated by dividing the number of abnormal landings by total attempted landings for all flies (n=3-15) for a single petri dish. Female flies were used for all heat assay genotypes except for htorΔE, where male flies were tested to maximize the sensitivity for detecting a potential phenotype.

The adult mechanosensitivity assay was adapted from the method described previously ^49^. Drosophila were collected as late-stage pupae and sequestered in standard food vials with 10-20 animals/vial and separated by sex. The animals were maintained at 25°C, 30-60% relative humidity with 12:12 light/dark cycle. Groups of 10-20 at 14 days post eclosion were allowed to acclimate to the environment for one hour. The animals were then transferred to empty vials and subjected to intense mechanosensory disturbance for 10 seconds input via Vortex Genie 2 on the highest speed. Recordings were collected until all animals recovered. The total time required to stand without restarting hyperkinetic movements was recorded for each fly by scorers blind to genotype. An animal was considered to be recovered after successful flight, walking up the side of the vial, or remaining stationary on its feet while grooming. Flies that did not respond to the stimulus were scored with a zero.

Graph creation and statistical test calculations were performed using R (4.2.2). For boxplots graphs, boxes represent 25^th^ and 75^th^ percentiles; whiskers represent 10^th^ and 90^th^ percentiles, and individual measurements are overplotted. Significance for average distance in vial assays determined using 2-tailed paired t-test. Difference in proportion of abnormal landings and hyperkinetic events calculated using 2-tailed unpaired t-test. The increase of total landings over time was calculated via 2-tailed unpaired t-test between baseline (30-40 seconds) versus other time bins. Gwet AC1 was used to test scorer agreement and to determine when a scorer had finished training using the R irrCAC package. For Gwet AC1 calculation, contingency tables were created between scorers across trials, using the number of all landings to identify agreement for a binary abnormal/normal classification.

### Immunohistochemistry and imaging at the neuromuscular junction

Imaging of neuromuscular junction of 3^rd^ instar wandering larva performed as previously described ^34,72^. Briefly, larval filet was dissected in HL-3 Ca++-free saline before fixation in Ca++ free 4% PFA. Washes performed with PBS with or without 0.1% Triton-X. Blocking before and during primary and secondary antibody steps using PBST with 5% normal goat serum, 2% BSA. Antibodies used in study listed in **Supplemental Table 2**. Primary antibody 1:400 M α DLG (4F3) incubated overnight at 4°C. DLG DSHB hybridoma monocolonal antibodies deposited by Goodman, C. Secondary antibodies 1:400 G α R Cy3 (ThermoFisher) incubated for 1.5 hours at RT. Neuronal membranes visualized with 1:100 Gα-HRP-Alexa 647 (Jackson ImmunoResearch) added with secondary antibody; HRP=horseradish peroxidase which recognizes neuronal membranes in Drosophila. 1:300 phallodin-488 in PBS (Molecular Probes) added as the final wash before mounting.

Imaging performed with 710 Zeiss confocal using 1.0 µm Z-stack at 63x of the 1b 6/7 muscle in the A3 abdominal segment. Image parameters (laser gain and intensity, resolution, zoom) were kept constant between images of the same session, alternating between control and mutants. Maximum intensity projections were created for a given field of view (FOV) using all parts of the Z-stack that included the axon terminal (defined by HRP staining) for all channels. Projections converted to greyscale from RGB in Photoshop (Adobe) without adjustments to pixel intensity. Non 6/7 1b HRP staining was identified based on shape/size/HRP staining intensity and manually removed.

Images analyzed using ImageJ software v. 1.50i (National Institutes of Health) by scorers blind to genotype. Images from HRP channel converted to black and white using threshold feature with the threshold held uniform for images within an imaging session. NMJ area measured by drawing around area encompassed by both muscles and neuron in the FOV. Orthogonal projections converted to 8-bit black and white tiffs without thresholding. Boutons were counted manually from 63x images, restricted to the same areas quantified for neuron area. Statistics and graphs created using R (4.2.2). For boxplots graphs, boxes represent 25^th^ and 75^th^ percentiles, whiskers represent 10^th^ and 90^th^ percentiles, and individual measurements are indicated with points.

### Treatment with ISRIB

We modified a previously described protocol for feeding ISRIB to developing larva ^73^. ISRIB was dissolved in a stock solution with DMSO and mixed with warmed fly food to a final concentration of 500 nM ISRIB and 0.25% bromophenol blue. Successful uptake of drug was confirmed by examining larval gut for bromophenol blue. Control animals were reared on an equivalent final concentration of DMSO (0.3%) and 0.25% bromophenol blue.

For survival, *Drosophila* were collected as late-stage pupae, separated by sex, and sequestered in standard food vials with ∼20 animals/vial. The number of alive animals were monitored daily, with animals transferred to fresh food once per week. Survival library in R (4.1.0) was used to generate Kaplan-Meier plots and the log-rank test statistic.

### Resource Availability Statements

This study did not generate new unique reagents or original code. All datasets available upon reasonable request.

## Notes

### Competing Interest Statement

The authors have declared no competing interest.

