## Supplemental materials for "eIF2α phosphorylation evokes dystonia-like movements with D2-receptor and cholinergic origin and abnormal neuronal connectivity"

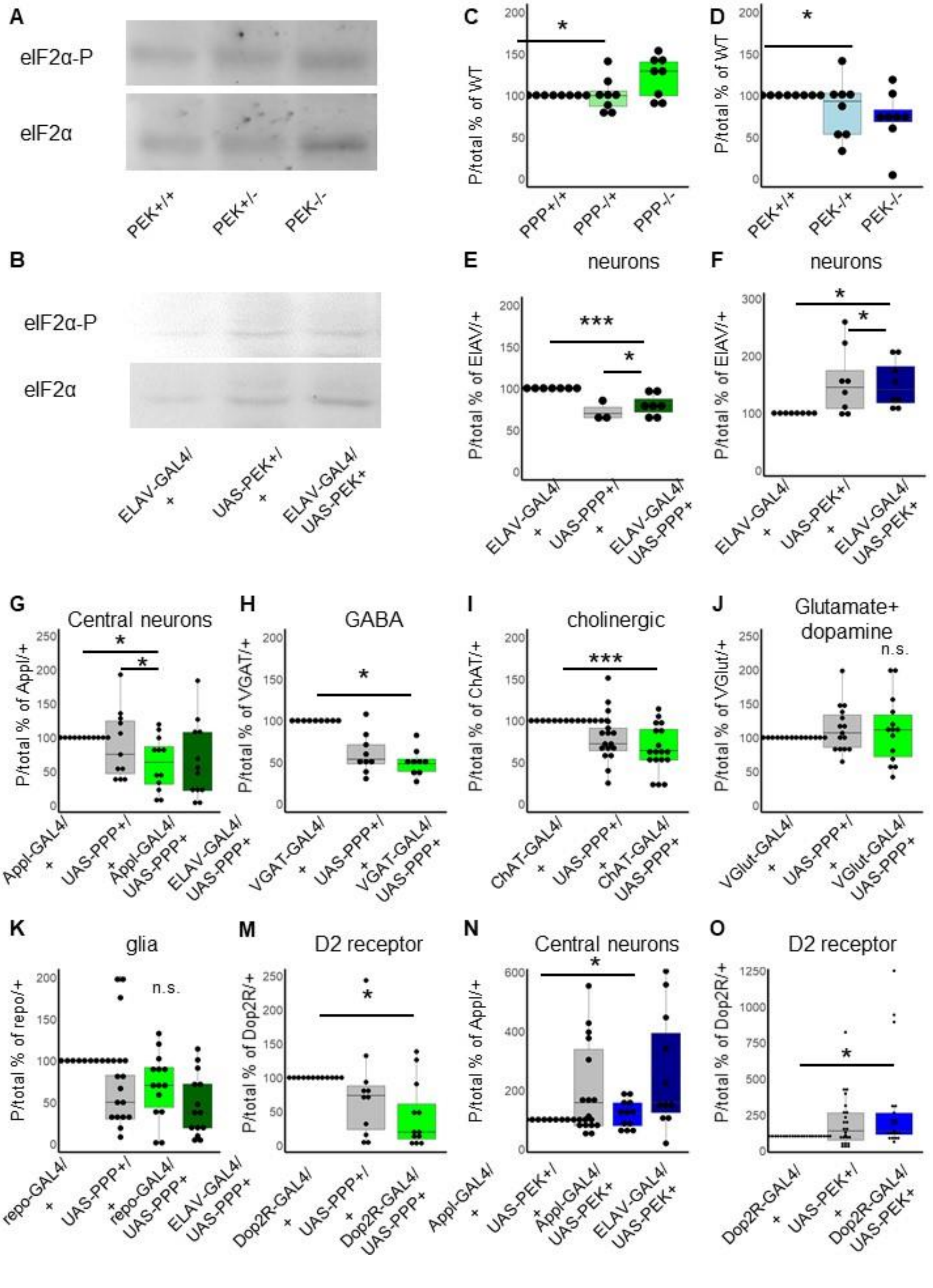

A
eIF2α-P
eIF2α
PEK+/+
PEK+/-
PEK-/-
B
eIF2α-P
eIF2α
ELAV-GAL4/ +
UAS-PEK+/ +
ELAV-GAL4/ UAS-PEK+
C
P/total % of WT
PPP+/+
PPP-/+
PPP-/-
D
P/total % of WT
PEK+/+
PEK-/+
PEK-/-
E
neurons
P/total % of ElAV/+
ELAV-GAL4/ +
UAS-PPP+/ +
ELAV-GAL4/ UAS-PPP+
F
neurons
P/total % of ElAV/+
ELAV-GAL4/ +
UAS-PEK+/ +
ELAV-GAL4/ UAS-PEK+
G
Central neurons
P/total % of Appl/+
Appl-GAL4/ +
UAS-PPP+/ +
Appl-GAL4/ UAS-PPP+
ELAV-GAL4/ UAS-PPP+
H
GABA
P/total % of VGAT/+
VGAT-GAL4/ +
UAS-PPP+/ +
VGAT-GAL4/ UAS-PPP+
I
cholinergic
P/total % of ChAT/+
ChAT-GAL4/ +
UAS-PPP+/ +
ChAT-GAL4/ UAS-PPP+
J
Glutamate+ dopamine
n.s.
P/total % of VGlut/+
VGlut-GAL4/ +
UAS-PPP+/ +
VGlut-GAL4/ UAS-PPP+
K
glia
n.s.
P/total % of repo/+
repo-GAL4/ +
UAS-PPP+/ +
repo-GAL4/ UAS-PPP+
ELAV-GAL4/ UAS-PPP+
M
D2 receptor
P/total % of Dop2R/+
Dop2R-GAL4/ +
UAS-PPP+/ +
Dop2R-GAL4/ UAS-PPP+
N
Central neurons
P/total % of Appl/+
Appl-GAL4/ +
UAS-PEK+/ +
Appl-GAL4/ UAS-PEK+
ELAV-GAL4/ UAS-PEK+
O
D2 receptor
P/total % of Dop2R/+
Dop2R-GAL4/ +
UAS-PEK+/ +
Dop2R-GAL4/ UAS-PEK+

Supplemental Figure 1: Alterations to eIF2a-P confirmed by western blotting. Green represents alterations to the PPP151R phosphatase (PPP), which removes phosphorylation from eIF2α (eIF2α-P). Blue represents the PERK kinase (PEK) which phosphorylates eIF2α-P. A-B, sample images of western blots. C-D. Quantification of ratios of phosphorylated/total eIF2α normalized to homozygous genetic background line (yw) for whole-animals collected at 14 days post-eclosion with loss-of-function variants in PPP (C) and PEK (D). Decreasing PPP increases ratios of eIF2α-P and decreasing PEK decreases ratios of eIF2α-P. E-O Quantification of ratios of phosphorylated/total eIF2α normalized to Gal4/+ heterozygous control taken from adult heads 14 days post-eclosion. E-F. Neuron-specific overexpression of PPP increases eIF2α-P (E) and PEK decreases eIF2α-P (F). G-M Overexpression of PPP in Gal4 lines decreases eIF2α-P and reaches significance in central (G), GABA (H), cholinergic (I), and D2 receptor (M) neurons. Non-significance is thought to be due to small anatomical regions of overexpression. N-O. Overexpression of PEK in Gal-4 lines significantly increases eIF2α-P in central (N) and D2-receptor (O) containing neurons. Dop1R n.s. (not shown).

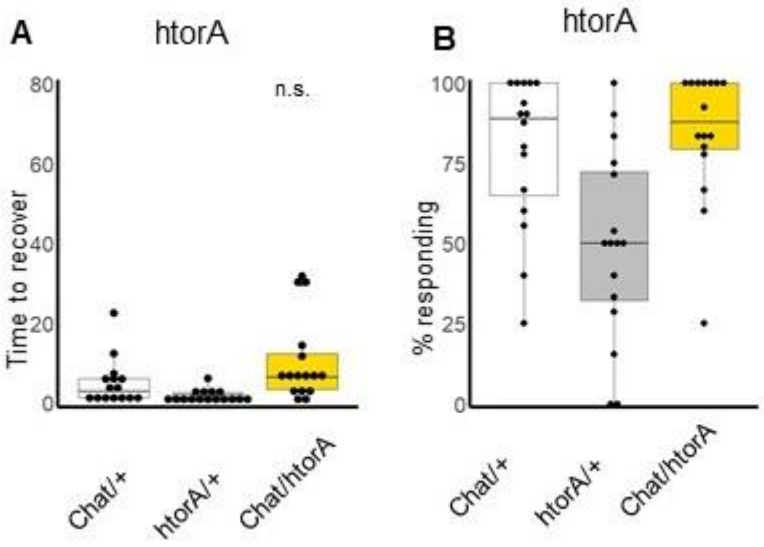


Supplemental Figure 2. Expressing a wildtype DYT1 allele, htorA, does not increase sensitivity to mechanostimulation-evoked dyskinetic movements compared to controls. A. No difference in time to recover. B. No difference in percent responders.

| Genotype abbreviation | Localization/timing | Genotype | Source |
|---|---|---|---|
| w1118 | whole animal | $w^{1118}$<br>crossed with Canton S and backcrossed for 12 generation | BDSC 3605 |
| y w | whole animal | y w | BDSC 6598 |
| CantonS | whole animal | wild-type (+) | BDSC 64349 |
| PPP- | whole animal | yw; P[w+]PPP1R15$^{G18907}$ 2$^{nd}$<br>5’ insertion<br>(UAS mis-expression element) | BDSC 26945 |
| PEK- | whole animal | yw; PEK$^{EY09578}$ 3$^{rd}$<br>5’ insertion<br>(UAS mis-expression element) | BDSC 17582 |
| ELAV | Post mitotic neurons, embryonic-adult | w*; P[GAL4-elav.L] 3$^{rd}$ | BDSC 8760 |
| Appl | Neurons, central brain> VNC>salivary glands+fat body. higher in males. Larva-adult. | P[Appl-Gal4.G1a]1, y w* 1$^{st}$ | BDSC 32040 |
| VGlut | Synaptic vesicle transporter in terminals of glutamatergic and dopaminergic neurons | TI[2A-GAL4]VGlut2A-GAL4/CyO 2$^{nd}$ | BDSC 84697 |
| Dop1R | Dopamine receptor GPCR; gamma Kenyon cell, adult MB, optic lobe | $w^{1118}$; P[GMR72B08-GAL4] attP2 3$^{rd}$ | BDSC 46669 |
| Dop2R | Dopamine receptor GPCR; nervous, digestive, excretory systems. Pre 3$^{rd}$ instar larva expression. Higher in males. | w* TI[2A-GAL4]Dop2R2A-GAL4 1$^{st}$ | BDSC 84628 |
| repo | glia. Embryonic-adult | $w^{1118}$; P[GAL4]repo/TM3, Sb 3$^{rd}$ | BDSC 7415 |
| VGat | GABA vesicle transport; inhibitory interneurons, embryonic-adult | w*; P[VGAT-GAL4.F] 3$^{rd}$ | BDSC 58409 |
| ChAT | Choline acetyltransferase; excitatory neurons, embryonic-adult | TI[2A-GAL4]ChAT2A-GAL4/TM3, Sb 3$^{rd}$ | BDSC 84618 |
| OK6 | motor neurons, embryonic-adult | P[GawB]OK6 2$^{nd}$ | BDSC 64199 |
| OK107 | MB neurons + neuroblasts, larval-adult | w; P[eyOK107/In(4)ci$^{D}$, pan$^{cID}$, sp$^{spa\text{-}pol}$ 4$^{th}$ | BDSC 854 |

| | | | |
|---|---|---|---|
| PPP+ | UAS-driven (Gal4 cell-specific) | M[UAS-PPP1R15.ORF.3xHA.GW]ZH-86Fb 3rd | Fly ORF F003018 |
| PEK+ | UAS-driven (Gal4 cell-specific) | $w^{1118}$; P[UAS-PEK.M]3/TM6B, Tb 3rd | BDSC 76248 |
| ATF4+ | UAS-driven (Gal4 cell-specific) | M[UAS-crc.ORF.3xHA.GW]ZH-86Fb 3rd | Fly ORF F000106 |
| htorΔE | UAS-driven (Gal4 cell-specific) | w-; P[w+, UAS-htorADeltaE]#24 2nd | Naoto Ito (Wakabayashi-Ito et al., 2015) |
| htorA | UAS-driven (Gal4 cell-specific) | w-; P[w+, UAS-htorA]#8 2nd | Naoto Ito (Wakabayashi-Ito et al., 2015) |
| $shibire^{ts\text{-}1}$ | UAS-driven (Gal4 cell-specific) | w*; P[UAS-shits1.K]3 | BDSC 442222 |

Supplemental table 1: Drosophila lines used in studies. BDSC=Bloomington Drosophila Stock Center. VNC=ventral nerve cord (spinal cord analogue). MB=Mushroom bodies. GPCR=G-coupled protein receptor.

| Antibody/stain | Species | Concentration | Application | Company | Lot(s) # |
| --- | --- | --- | --- | --- | --- |
| DLG | M | 1:400 | NMJ IHC | DSHB 4F3 | 2/18/16 |
| Anti-mouse Cy3 | G | 1:400 | NMJ IHC | ThermoFisher | A10521 1425618 |
| Anti-HRP 647 | G | 1:100 | NMJ IHC | Jackson Immuno Research | 126323 |
| Phalloidin 488 | - | 1:300 | NMJ IHC | Molecular Probes | 1903540 2160010 |
| Beta actin | M | 1:2000 | WB | Abcam 8224 | GR14272-3 |
| Beta tubulin | R | 1:2000 | WB | Abcam 6046 | GR3376491-1 |
| Phosphor-S51 eIF2S | R | 1:1000 | WB | Cell Signaling 3597 | 12 |
| eIF2S1 | R | 1:500 | WB | Abcam 26197 | GR3183673-1 GR3325905 |
| Rabbit ECL | G | 1:10000 | WB | GE healthcare NA931 | 17473046 |
| Mouse ECL | G | 1:10000 | WB | GE Healthcare NA934 | 17170583 |

Supplemental Table 2: Antibodies used in studies. HRP=horseradish peroxidase (recognizes nirvana2 neuronal surface protein in Drosophila), ECL= Enhanced Chemiluminescence, M=mouse, G=goat, R=rabbit. NMJ IHC=neuromuscular junction immunohistochemistry, WB=western blot

| Software | Library/ function | Company | Source |
| --- | --- | --- | --- |
| R/R studio (v. 4.2.2) | Survival, tidyverse, irrCAC | The R project for Statistical Computing | https://cran.r-project.org/bin/windows/base/ |
| ImageJ software v. 1.50i | measure distance traveled and NMJ area | National Institutes of Health | https://imagej.nih.gov/ij/download.html |
| Photoshop | image stitching | Adobe | https://www.adobe.com/products/photoshop.html |
| Premier Pro/Rush | frame-by-frame video analysis | Adobe | https://www.adobe.com/products/premiere.html |
| Image studio Lite | western signal quantification | LiCor | https://www.licor.com/bio/image-studio-lite/ |
| Zen Black/blue | confocal imaging and projections | Zeiss | https://www.zeiss.com/microscopy/en/products/software/zeiss-zen-lite.html |

Supplemental Table 3: Software used in studies and sources.
